# High-throughput microcolony growth analysis from suboptimal low-magnification micrographs

**DOI:** 10.1101/253724

**Authors:** Yevgeniy Plavskin, Shuang Li, Hyun Jung, Federica M. O. Sartori, Cassandra Buzby, Heiko Müller, Naomi Ziv, Sasha F. Levy, Mark L. Siegal

## Abstract

New technological advances have enabled high-throughput phenotyping at the single-cell level, yet analyzing the large amount of data generated by high throughput phenotyping experiments automatically and accurately is a considerable challenge. Here we introduce Processing Images Easily (PIE), software that automatically tracks growth of microbial colonies in low-magnification brightfield images by combining adaptive object-center recognition with gradient-based object-outline recognition. PIE recognizes colony outlines very robustly and accurately across a wide range of image brightnesses, focal depths, and organisms. Beyond accurate colony recognition, PIE is designed to easily integrate with complex experiments, allowing colony tracking across multiple experimental phases and classification based on fluorescence intensity. We show that PIE can be used to accurately measure the growth rates of large numbers (>90,000) of bacterial or yeast microcolonies in a single-time-lapse experiment, allowing calculation of population-wide growth properties. Finally, PIE is able to track individual colonies across multiple experimental phases, measuring both growth and fluorescence properties of the microcolonies.

**Author Summary:** High-throughput microscopy has enabled automated collection of large amounts of growth and gene-expression data in microbes. Computational methods that can precisely recognize and track organisms in images are essential to performing measurements at scale using automated microscopy. We have developed PIE, software that automatically recognizes microbial colonies in microscopy images, tracks them in imaging time-series, and performs measurements of growth and, potentially, gene expression. PIE is highly effective on low-resolution images, outperforming current state-of-the-art approaches in both speed and accuracy, and works well in microbes of varying shapes and sizes. In addition, PIE allows tracking microcolonies across arbitrary sequences of experimental phases, each collecting data in different modalities. We show that PIE allows measurement of growth and fluorescence properties in tens of thousands of microbial colonies in a single experiment, and that in turn the scale of these measurements can lead to important insights about interindividual differences in growth and stress response. PIE is available as a Python package (https://doi.org/10.5281/zenodo.4987127) with documentation currently at https://pie-image.readthedocs.io/; users can also run analysis on individual images or time-series without the need to install PIE by using our web application, currently available at http://pie.hpc.nyu.edu/.

## Introduction

The development of new technologies for high-throughput phenotyping has allowed increasingly precise measurements of many traits through vastly increased sample sizes, as well as the ability to explore previously intractable traits, such as the extent to which genetically identical individuals raised in the same environment vary [1]. Automated microscopy of immobilized organisms can produce large amounts of data on cell shape, internal structure, and growth patterns [2–4]. However, subsequent data analysis often presents a significant challenge, as machine-vision approaches must be applied to automatically extract useful data from the large number of images created.

One particularly important phenotype is the rate of cell growth. In multicellular organisms, cell growth is key to normal development and to disease processes such as tumorigenesis. In microbes, growth rate is closely related to fitness. High-throughput imaging allows population growth rate to be measured as a function of the growth of individual colonies [5] or microcolonies (e.g. [2,6–9]) or the growth and division times of individual cells (e.g. [10–12]) rather than a single population-wide average. Individual measurements, in turn, allow exploration of important properties of the growth-rate distribution within a population, such as dispersion and skewness. For example, in the budding yeast *Saccharomyces cerevisiae*, the distribution of microcolony growth rates differs among strains and tends to have a heavy tail of slower-growing microcolonies [2,8,13]. Heterogeneity is especially well-measured in microcolony approaches because of the small number of cells per colony and the large scale of experiments possible.

To automatically measure microcolony growth rates from microscope-based images, two computational problems need to be solved: recognition of colonies and tracking of colonies through subsequent time points. Brightfield microscopy requires the least specialized equipment (unlike e.g. phase-contrast microscopy) and no specially marked strains (unlike fluorescent microscopy), but segmentation of cells and microcolonies can often be most difficult from brightfield images. CellProfiler identifies, segments, and tracks cells in images (Carpenter et al., 2006). However, optimization of object-recognition algorithm parameters for CellProfiler can make this approach time consuming to customize and test for each application. Dedicated software exists that addresses the problem of cell recognition in brightfield images (e.g. YeastSpotter for *S. cerevisae*, Cellpose for diverse cell types [14,15]), but these applications do not allow for tracking microcolonies across time, and were trained primarily on high-resolution micrographs, potentially limiting their effectiveness on lower-resolution images.

We aimed to develop a method to perform automated recognition and tracking of brightfield microbial microcolonies. Such a method must be robust to imaging variation both across and within experiments, needs to allow flexibility in experimental setup, and ideally would be easily adaptable to different species. Here, we introduce Processing Images Easily (PIE), an algorithm for tracking growing microbial microcolonies in low-resolution brightfield and phase-contrast images. Our image-processing procedure combines adaptive object-center recognition based on relative brightness with gradient-based object-outline recognition, resulting in a high level of robustness to image brightness and focal depth while minimizing the number of user-supplied parameters. PIE’s colony tracking is both more robust and >10x faster than previously published methods, allowing for fast, accurate colony tracking from large-scale microscopy experiments.

The colony-tracking procedure then joins tracked colonies across sequential time points, and calculates growth rates and pre-growth lag times, as well as simultaneous tracking of other colony properties (e.g. fluorescence) over time. We show that PIE can be used to collect data on the distributions of growth parameters across entire populations of cells in microbial species with diverse cell shapes and sizes. Finally, we provide an example of using PIE to track microcolony growth and fluorescence across multiple phases of an experiment, allowing properties of tens of thousands of individual microcolonies to be measured before and after administration of a stress.

## Results

### Software implementation

#### Procedure Overview

The growth-rate measurement algorithm described here consists of two stages: colony recognition, followed by colony tracking through time series (or sequential phases). Key colony parameters for each time point are recorded for growth-rate estimation and downstream analyses.

Colony recognition is divided into three substages (**Fig 1**). First, an automated threshold on relative image brightness identifies ‘cell center’ objects within an image. Next, the image is split into ‘gradient change’ objects whose edges coincide with gradient direction changes in the image. Finally, each colony is identified as the union of contiguous gradient-change objects overlapping a cell center. An optional recursive ‘cleanup’ step removes spurious gradient-change objects that result from brightness fluctuations in the backgrounds of some images.

**Fig 1.**
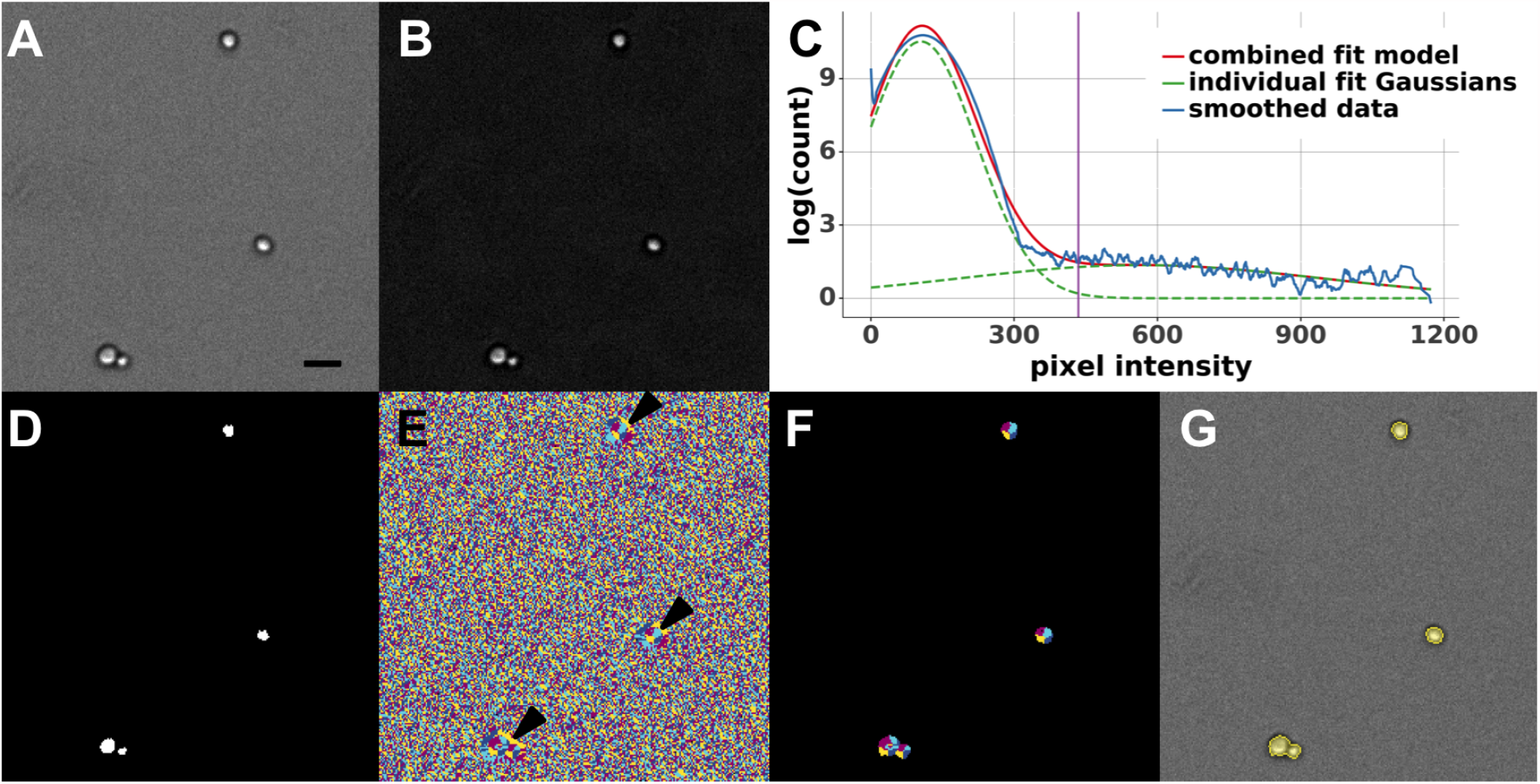
Steps of colony recognition by Processing Images Easily (PIE). (A) Raw brightfield image. To display each colony object clearly, only a representative part of the image is shown; scale bar is 10 µm. (B) Top-hat filtered image. (C) Smoothed log-frequency histogram of the top-hat image with its two-Gaussian fit. Vertical axis is the density of the log frequencies of pixel intensities. Blue solid curve shows the smoothed log-frequency histogram of the top-hat image; green dashed curves are the two Gaussian fits; red solid curve is the fit combining the two Gaussians; purple vertical line is the automatically calculated threshold for cell centers in the top-hat image. (D) Cell centers after thresholding the top-hat image. (E) Combination of four gradient-changes of the raw brightfield image. Four gradient-changes are: decreasing in x (left-to-right) and increasing in y (top-to-bottom) (quadrant I, light blue); increasing in x and increasing in y (quadrant II, purple); increasing in x and decreasing in y (quadrant III, yellow); and decreasing in x and decreasing in y (quadrant IV, dark blue). (F) Gradient-change objects from (E) that overlap with cell centers (arrowheads) from (D). Color code is the same as in (E). (G) Segmentation (yellow shading and lines) of final colony objects recognized by PIE overlaid on raw brightfield image.

During the colony tracking stage (**Fig 2**), sequential pairs of images are aligned, and colonies are tracked forward through time to identify which colony or colonies in a preceding time point’s image spatially overlap with each colony in the current time point’s image [2]. Colony fusion and splitting events are identified. In the case of fusion, tracking ends for both fused colonies. In the case of splitting, colony areas are calculated by including both the major colony and the components that have split off. After tracking colony identities between images, colony areas and other properties called for by the experiment (e.g. colony fluorescence level in corresponding fluorescent-channel images) are then recorded for each time point.

**Figure 2.**
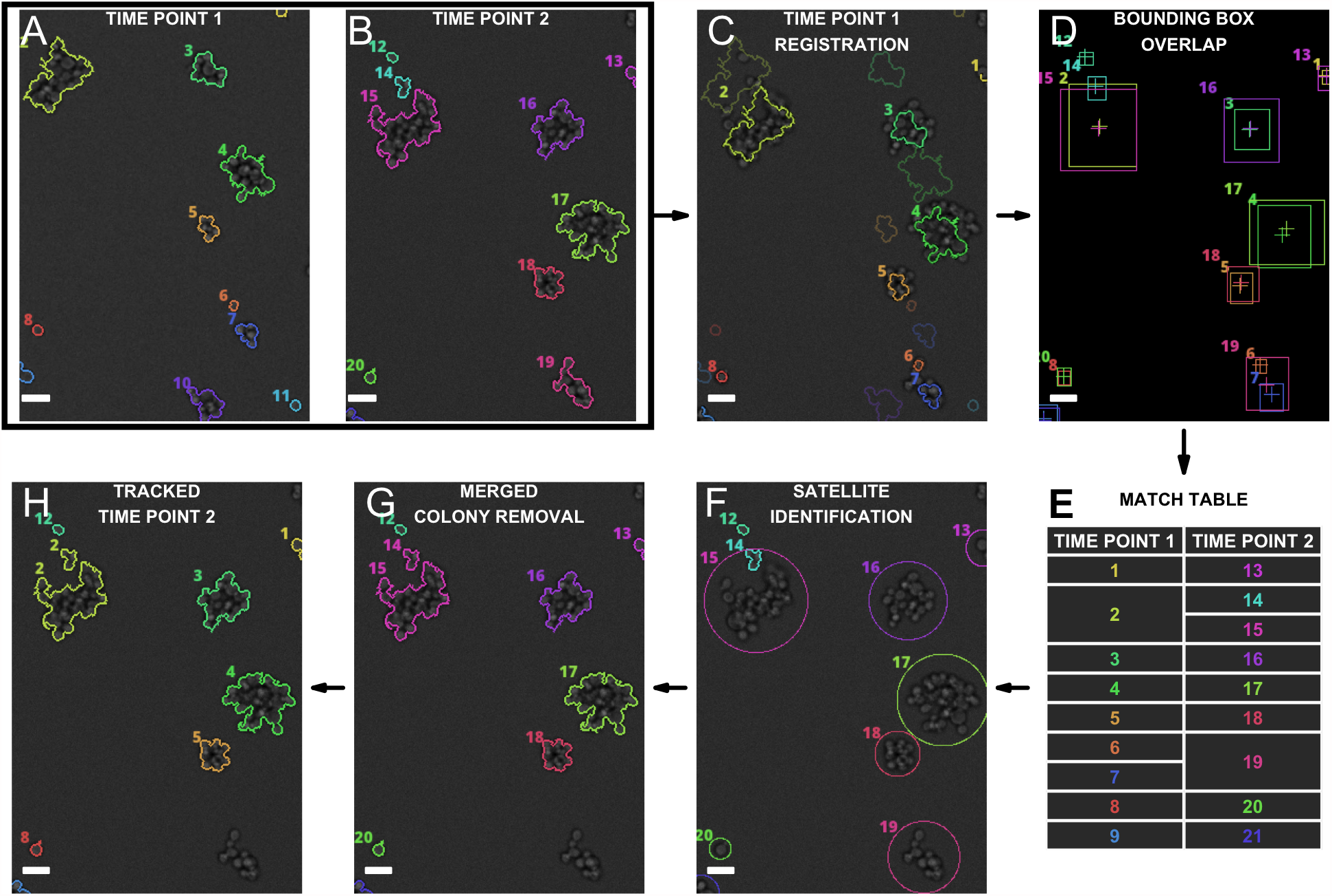
Colony tracking algorithm. (A-B) Two images of the same position from sequential time points, time point 1 and time point 2 (tp1 and tp2), with numbered recognized colonies in each. Note that the colonies are not perfectly aligned between these two images (top left). (C) The optimal affine transformation (translation + rotation) between the images in tp1 and tp2 is found, and tp1 colony positions are transformed to align them with tp2 colonies (shown in the image are tp1 colony outlines and centers overlaid on tp2 image; original positions are in faint colors, tp2-registered positions are in bright colors.) (D) Overlaps between each colony’s bounding boxes from tp1 and tp2 are calculated; colony centers indicated as crosses. (E) Matches are tabulated for any colony whose center in one time point is within the bounding box of a colony from the other time point. Note the ‘split’ event, in which a colony from tp1 (colony 2) matches two colonies in tp2 (colonies 14, 15); of these, only the largest will continue to be followed as the “main” colony. Also note the ‘merge’ event, in which two colonies in tp1 (colonies 6, 7) match a single colony in tp2 (19). Note that some colonies may remain unmatched (e.g. colony 12 in tp2, top left). (F) Any colonies with no match to the previous timepoint (including smaller colonies in split events) are assigned as ‘satellites’ to their larger ‘parent’ colony if their center falls within a size-dependent radius around a matched colony, as detailed in the text (here, colony 14 becomes counted as part of colony 15 for purposes of future tracking and area calculation). (G) Any colonies resulting from a merge (and their satellites) are removed from tp2 (here, colony 19 has been removed) (H) Identities of any matched colonies are propagated from tp1 to tp2, and the process is repeated between tp2 and the subsequent time point. Scale bar is 10 µm.

#### Cell center recognition

Cell centers are recognized by identifying an appropriate intensity threshold on a locally background-corrected image. An original image (**Fig 1A**) is first subjected to a top-hat filter [16] to correct for uneven illumination, thereby reducing background and making it more uniform while preserving the brightest parts of the image, which correspond to cell bodies (**Fig 1B**). The appropriate threshold for cell-center recognition is identified using a smoothed histogram representing the log frequencies of pixel intensities of the top-hat-filtered image. Where possible, the histogram bin centers are set to the unique pixel intensity values in the image. However, if the number of unique pixel intensity values exceeds a heuristic image size-dependent value, evenly spaced bins are used. In this case, a bin number is selected such that there are no peaks in the log frequency histogram autocorrelation (this autocorrelation check is necessary to avoid the poor and discontinuous binning that often results from using evenly spaced bins on images). The log-frequency histogram is then smoothed by applying the Savitzky-Golay filter [17], which fits successive windows of adjacent data points to low-degree polynomials. This step eliminates small spiky peaks of the histogram, which would otherwise introduce error in threshold selection.

In a typical brightfield image of cells, the smoothed log-frequency histogram of the top-hat image resembles a truncated mixture of two Gaussian curves (**Fig 1C**): one Gaussian curve corresponds to background pixels, which have lower pixel intensities, and one corresponds to bright cell bodies, which have higher pixel intensities. A good threshold to capture cell centers will sit in between the peaks of the two Gaussian curves. (In rare cases, phase-contrast images require three Gaussians to be fit, and users can select to perform thresholding this way; in that case, the ideal threshold rests between the peaks of the first and second Gaussian.) Because cell bodies appear as an intensity gradient, with the center being the brightest, and because cell outline recognition does not depend on the exact outline of the cell center detected here, a range of thresholds will produce satisfactory results.

Automated threshold identification is implemented on the smoothed log-frequency histogram of the top-hat image by one of two methods: by either fitting a mixture of Gaussians or applying a sliding circle to identify the peak-separating threshold. Our algorithm first attempts to fit a mixture of Gaussians to the smoothed log-frequency histogram of the top-hat image (**Fig 1C, red curve**). The smoothed log-frequency histogram of the top-hat image is first fit, by non-linear least squares, to a two- (or three-)Gaussian mixture model with one Gaussian constrained to have mean greater than 0. A number of log-frequency histogram quality controls and the starting parameter values of the distribution fitting are important to achieve this initial good fit, and are described in detail in the PIE code.

In all test images tried, the approach described above resulted in a good fit of the curve to the smoothed log-frequency histogram (adjusted R^2^ > 0.85), producing an acceptable threshold value. However, it is possible that this method would fail to produce a satisfactory fit in images in which the pixel intensity distribution differed drastically from our test images. To prevent thresholding failure in unforeseen imaging conditions, we have built in alternatives to capture the pixel distributions possible with a two-Gaussian fit. As described in **S1-S2 Figs**, these alternate approaches are tried sequentially by PIE until a satisfactory fit to the smoothed log-frequency histogram is found. If all the Gaussian fitting methods fail to produce a satisfactory fit, a more time-consuming sliding-circle approach is used to identify a threshold separating the peak representing background pixels from the one representing bright cell center pixels.

To identify cell centers, the determined threshold is applied to the top-hat-filtered image (**Fig 1D**), resulting in a binary mask that mostly corresponds to the central regions of cells.

#### Colony outline recognition

To detect colony outlines, PIE uses a gradient-based approach that is robust to differences in image brightness and, to some degree, image focal position. We first divide images into objects whose outlines correspond to shifts in gradient direction. Objects that do not overlap the cell centers identified above are then removed, leaving microcolony objects.

To split the image based on gradient direction, a Sobel gradient filter [16] is applied to the image separately in the x and y directions; a threshold of 0 is then applied to the negative and positive versions of each of the gradient images, and pairs of resulting images are multiplied elementwise. The result is a set of four binary images, with objects representing each possible combination of gradient directions in each image: increasing in x and increasing in y, increasing in x and decreasing in y, decreasing in x and increasing in y, and decreasing in x and decreasing in y. For an object with a bright center and a dark outline, such as a slightly defocused image of a cell, this procedure results in objects in four quadrants resembling pie pieces (**Fig 1E**). To identify these pie-piece-shaped objects belonging to cells, any objects not overlapping a cell center identified in the previous section are removed (**Fig 1F**). This leaves only those objects that belong to cells. Contiguous “pie piece” quadrants are then merged into colony objects; as a result, any cells that are touching are considered to be part of a single colony (such as the mother cell and bud cell on the bottom left of **Fig 1F**). Internal holes smaller than an optional user-specified parameter are filled to create the final colony objects (**Fig 1G**).

We find that in some imaging routines, microcolonies are surrounded by bright areas of background. The cell center-finding algorithm described above identifies the cell-adjacent bright background regions as cell centers, and as a result, pie pieces in the background are incorrectly identified as microcolonies. Our procedure allows for an optional recursive ‘*cleanup*’ procedure for sets of images in which this problem occurs frequently (**S3A Fig**). This cleanup is optional because it substantially extends runtimes and could result in inconsistent tracking of small cells, especially at early time points (**S3B Fig**).

The cleanup procedure has two steps. The first step is based on the observation that, unlike most true pie pieces corresponding to cells, the spurious background pie pieces are often irregularly shaped and have long, exposed edges that are not in contact with neighboring pie pieces. The cleanup procedure takes advantage of this observation by removing any pie pieces for which the proportion of the perimeter pixels not in contact with another center-overlapping pie piece is less than 0.75 (although this value can be adjusted by the user). The second step in the cleanup procedure takes advantage of the fact that, for an object with a bright center and dark edges (such as a cell), the gradient should radiate from the center out, meaning that the pie pieces that compose the cell should be in a predictable arrangement. Specifically, a negative x gradient and positive y gradient in the upper right quadrant should neighbor a negative x gradient and negative y gradient in the bottom right quadrant, and a positive x gradient and positive y gradient in the upper left quadrant, etc (**Fig 1F**). Any center-overlapping pie pieces whose neighbors do not have the expected gradient direction are removed. The two cleanup steps are performed recursively, until no more pieces are removed.

Finally, rare failures in thresholding (for example, in images with backgrounds consisting of two distinct intensities) can result in identification of a very large number of spurious colony objects, causing incorrect colony identification and taking up significant computational resources to analyze. PIE therefore removes any images in which the number of detected colonies exceeds an optional user-defined parameter, treating those images as blank.

#### Fluorescence Measurements

For any fluorescent channels, PIE reports the mean, median, and upper quartile of fluorescence values for each colony, as well as for the local ‘background’ area around the colony. To calculate fluorescence inside the colony, the binary colony mask is eroded with a circular structure element with a radius of 2 to make the mask slightly smaller than the recognized colony area. The eroded mask is then used to retrieve fluorescence intensity statistics from each channel. The use of an eroded mask makes fluorescence measurements more robust to slight errors in colony-outline recognition and to inconsistencies in fluorescence at cell edges. Although this approach would be problematic for fluorescent markers of the cell wall or cell membrane, the underlying scripts can be modified to include the eroded border pixels in the fluorescence calculation. PIE also reports the local background fluorescence in each channel, measured within an area of the colony bounding box (the smallest rectangle containing the identified colony) extended by 5 pixels in all four directions but outside of the colony area itself, or of any other colony within the extended bounding box. The resulting background mask is eroded with a disk structure element with a radius of 2 for the same reason as the colony mask. Reporting both colony and background fluorescence intensity values in this way allows the user to perform local colony background correction.

#### Colony tracking through time and between experimental phases

Colony tracking is summarized in **Fig 2**. Colony identities and properties, such as area, centroid, pixel-coordinate list and bounding box are processed for all images at a given imaging position. Brute force matching of colony centroid coordinates in sequential images is used to identify the ‘match’ candidate centroid locations from one image to the next; only those matches where the best-matched centroid is < 0.75 times the distance to the second-best centroid are retained [18,19]. The resulting list of matching locations between sequential images is used to estimate a best-fit affine transform matrix. This matrix is then used to register the image at the first time point to the image at the subsequent time point, aligning the colony locations. This image registration is important because it allows colonies to continue to be tracked even if the imaging field shifts between two time points, which often happens as a result of microplate thermal expansion or when treatments are applied to the cells mid-experiment; however, users have the option of turning off image registration.

After image registration, colonies are connected between sequential images. If the x and y distances between the centroids of two colonies in the current and the subsequent image are less than half the current or the subsequent bounding box width and height, then the two colonies are considered connected between these two time points. A colony may be connected to more than one colony in the previous image, meaning multiple colonies have merged; or one colony may split into several pieces, often creating ‘satellites’. All colonies that are connected to at least one other colony in a subsequent time point are classified as connected objects.

Merging colonies are treated differently than splitting colonies. Upon colony merging, tracking stops; that is, only properties before merging are recorded. When a colony splits the largest colony in the later time point is selected as the tracked colony object. This selection helps sustain tracking if a cell or small number of cells breaks off from a larger colony. Untracked colonies (including the smaller fragments of a split colony) that are close to an existing tracked colony are declared ‘satellite’ colonies. In our experience, defining satellites as being closer than 70% of the tracked colony’s long axis length produces robust tracking [2,6–9]. Importantly, merges between a satellite and its tracked parent colony do not result in a cessation of tracking, which means that PIE can continue tracking colonies when cells split off and land nearby, or when small colony recognition errors separate a single cell from its parent colony at a single timepoint.

If a satellite colony is identified, its area is added to its corresponding tracked colony at that particular time point. All other properties, such as centroids and fluorescent intensities, are still determined only based on the corresponding tracked colony. For fluorescent intensities, satellites are not included because the risk of counting brightly autofluorescent debris as a satellite outweighs the benefit of having a small number of additional cells from which to measure a colony’s overall per-pixel fluorescence (e.g., by the mean, median or upper-quartile fluorescence). If the satellites of a split colony fail to be identified as such, a significant drop in apparent colony area over time might result. However, we apply an optional filter during the growth rate analysis step to remove colonies whose areas dip significantly in successive time points, thus eliminating any split colonies in which satellite assignment was not successful.

#### Growth rate and lag calculation and data filtration

Although the colony recognition in PIE is robust across a range of imaging conditions, colony tracking failures still occasionally occur with large amounts of data. As a result, we use user-defined parameters to perform a number of filtration steps before and after growth rate and lag calculation.

First, PIE removes any tracked objects outside a range of acceptable colony area sizes that can be defined by the user based on knowledge of the study organism and imaging conditions. This filter can help prevent debris from being considered a colony. To remove colonies poorly tracked by PIE, user-defined filters can also be applied based on the maximum allowed decrease in pixel number and the maximum allowed fold decrease and increase in area between subsequent time points.

Growth rates and lag durations in PIE are calculated using a sliding-window approach similar to the method previously described [9]. A user-defined parameter defines the number of time points to be used per growth-calculation window; a colony must be tracked at every time point within the window (i.e. time points in which the colony was filtered out for any of the reasons described above, or for which an image is missing, may not be part of the window). For each tracked colony, a linear model is fit to the log of colony areas as a function of time (in hours). The slope of this regression line is the growth rate for the window. The lag duration is estimated based on the intersection of the regression line with a horizontal line determined by the area of the microcolony at the first time point at which it is tracked; the time between this intersection point and the start of imaging is the lag for the window. Finally, the linear model’s coefficient of determination (R^2^) and the fold increase in area are recorded for each growth window.

One growth window is chosen as the one from which the colony’s growth rate and lag duration are recorded. A growth window is removed from consideration if it fails to pass a minimum optional user-defined fold increase in area (which can be used to remove dead cells and debris from analysis), or if its R^2^ value is below an optional user-defined threshold; we recommend care when applying this threshold, as it will result in exclusion of colonies with growth rates close to 0. For any colonies that still have multiple growth windows after these filters, the window corresponding to the largest growth rate is selected. This selection decreases the chance that reported growth rates and lags will be based on growth windows that include either the colony’s lag period or the period after which the colony ceases to grow in two dimensions, when the change in colony area no longer reflects the full growth rate.

To prevent tracking of cells that come loose from existing colonies (but are not close enough to the parent colony to be counted as ‘satellites’), any colonies that do not appear in the first time point are removed. In addition, any colonies that are not tracked for a user-defined minimum of timepoints are optionally removed; this allows users to require colonies be tracked for longer than the number of time points used to calculate growth rate, or can serve to prevent transitory tracked objects from later resulting in removal of nearby objects due to minimal distance restrictions (see below).

A final optional filter removes growth-rate measurement biases introduced by colony density. Fast-growing neighboring colonies are more likely to merge with each other early in the experiment (i.e. before sufficient time points are collected for a growth window) than slow-growing colonies, resulting in more slow-growing colonies being counted, especially when cells are densely plated [2]. PIE therefore removes any colonies whose centroid falls within the user-provided minimum distance of another colony’s centroid at any point during the colony’s growth.

#### Multiphase experiments

PIE allows users to perform experiments in which microcolonies are tracked over multiple experimental phases, for example before and after a heat shock [2,7]. Each phase is initially treated as a separate experiment, with microcolony properties (e.g. growth rates, lags, and fluorescence intensities) calculated independently; user-specified filtration parameters and imaging parameters (e.g. fluorescent channels) can all differ between experimental phases, although data must be collected from the same imaging x-y positions in all phases of the experiment. For each phase, PIE performs temporal tracking of each colony within the phase, and in addition tracks colonies from the final time point of each phase to the first time point of the subsequent phase (see “Colony tracking through time and between experimental phases”). Colonies are assigned both within-phase and cross-phase tracking IDs, and colony properties such as growth rate are reported for each colony in each phase in which it was tracked.

#### Fluorescence-based classification

PIE is also capable of supporting experiments in which multiple fluorescent protein-marked strains are co-cultured in a single well, allowing classification of colonies based on mean colony fluorescence intensity at a single time point after growth rate data are collected. Users can specify an experimental phase for classification that consists of fluorescent imaging and has a ‘linked’ growth-rate phase; the fluorescent images from the classification phase are used in conjunction with colony outlines detected in the linked phase to assign a fluorescence value to each colony in each imaged channel. Binary classification is then performed for all colonies in each fluorescence channel: after the brightest 0.01% of colonies are removed, PIE fits a bimodal logistic distribution to the colony fluorescence data. PIE next identifies two thresholds: a ‘background’ threshold, and a ‘fluorescent’ threshold; colonies below the background threshold are classified as fluorescence-negative, colonies above the fluorescent threshold are classified as fluorescence-positive, and colonies between the thresholds are not classified. Each threshold is selected in such a way that the proportion of colonies that are misidentified as ‘background’ or ‘fluorescent’ is ∼0.1%, assuming that colony fluorescence values are drawn from the two logistic distributions estimated earlier.

### Robust colony recognition across focal planes and illuminations

To determine how accurately PIE recognizes microbes in low-resolution microscopy images, we first benchmarked the accuracy of PIE recognition of *Saccharomyces cerevisiae* microcolonies against three other approaches: a ‘naive’ approach consisting of automated ‘Maximum Entropy’ thresholding followed by a Hough Transform in Fiji [20–22], and two machine learning-based approaches: YeastSpotter, which was developed specifically for the task of detecting yeast cells in microscopy images [14], and CellPose, a general cell segmentation algorithm [15]. YeastSpotter and CellPose have both been shown to perform well on high-resolution cell images. We sought to determine whether these off-the-shelf approaches could robustly detect cells in low-resolution images, and to compare their performance to PIE’s. Colony recognition by all four methods was compared to a manually traced reference colony mask. To allow for direct comparisons between PIE (which recognizes microcolonies) and YeastSpotter and CellPose (which recognize individual cells and did not successfully segment multiple cells in large microcolonies in tested images), we benchmarked these methods on a set of microcolonies that had recently started growing and consisted primarily of one or two cells.

In comparing these approaches, we are concerned with both the accuracy of a colony’s outline given that the colony has been recognized, as well as the accuracy of recognizing colony objects in the first place. We therefore use two measures commonly used in image recognition studies: Intersection over Union (IoU) (**Materials and Methods, Eqtn 1**), which provides an estimate of the accuracy of colony outline recognition; and the F_1_ score (**Materials and Methods, Eqtn 2**), which reports the accuracy of recognizing colony objects. An optimal method will maximize both IoU and the F_1_ score. We therefore assessed overall accuracy across a range of IoU thresholds, by eliminating colonies that did not surpass the threshold (those that overlapped insufficiently with the benchmarking reference) and then computing the F1 score for the remaining colonies.

Assessing the F_1_ score across a range of IoU cutoff values shows superior performance for PIE (**Fig 3A**). Although the Hough Transform-based and CellPose methods match more colonies than PIE at low IoU cutoff values, the number of matches swiftly decreases as the IoU requirement becomes more stringent. The reason for this decrease is apparent in **Fig 3C**, which shows recognition results for a small subset of colonies in our test image: both CellPose and the Hough Transform method fail to recognize the entire cell in the image, instead recognizing only a bright area near the cell center. YeastSpotter, on the other hand, tends to include additional area around the cell in its mask, and occasionally misses cells. PIE generally recognizes most colonies in the image with an accurate cell outline, as reflected in the high F_1_ score across a wide range of IoU values.

**Fig 3.**
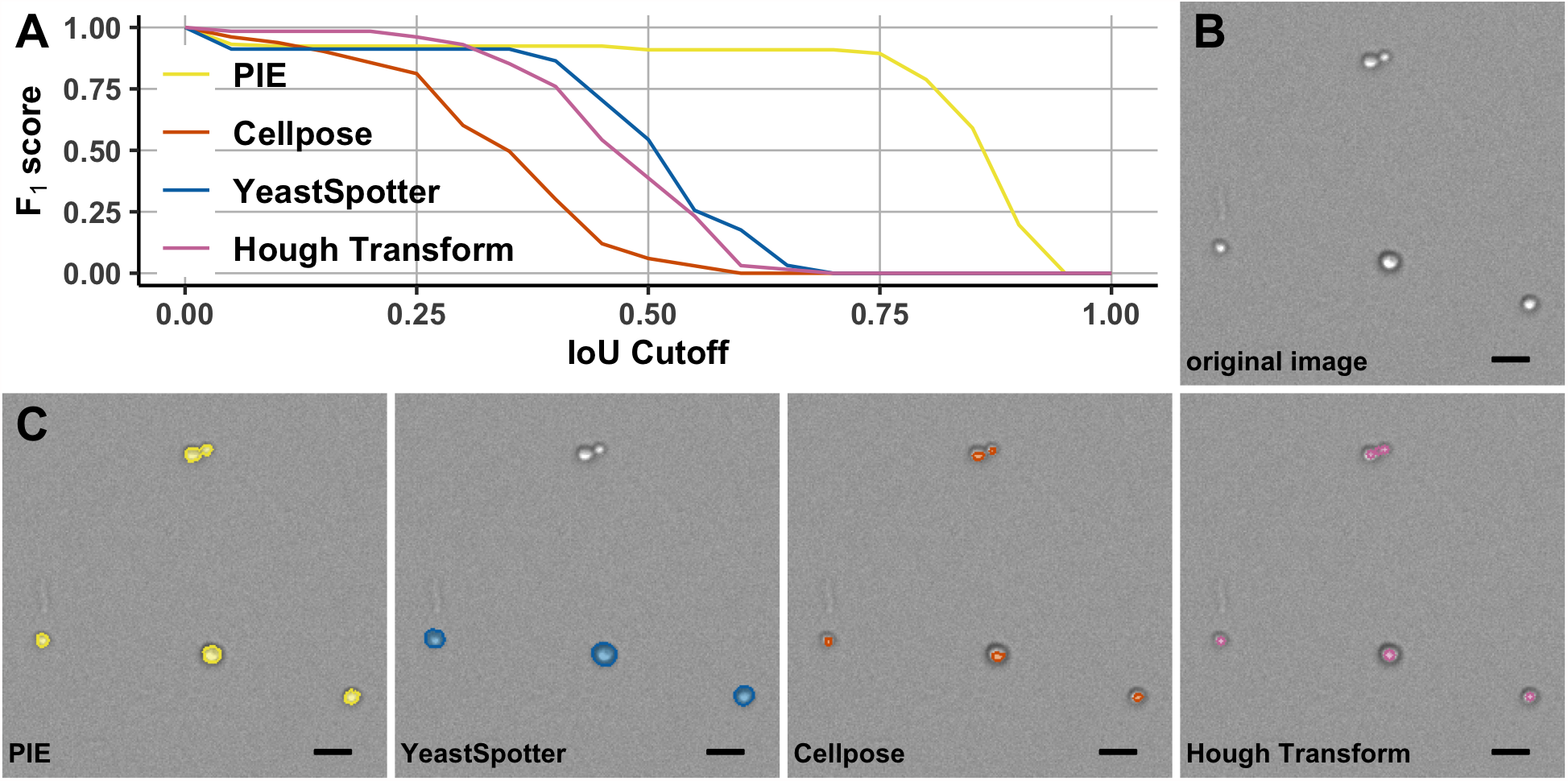
Benchmarking PIE performance on yeast cell recognition. (A) PIE achieves a high accuracy of microcolony recognition (F_1_ score) even at stringent requirements for accuracy of microcolony outline recognition (IoU); by contrast, the other methods only detect the majority of colonies with relaxed requirements for accuracy of microcolony outline recognition. (B) A subsection of the image used for benchmarking algorithms in (*A*) and (*C*). (C) Performance of the four algorithms overlaid on the image shown in (*B*). PIE accurately identified microcolony outlines; YeastSpotter failed to identify some colonies, and generally determined the cell outline to be outside of the bounds of the microcolony, resulting in low IoU values for many cells. While CellPose and Hough Transform methods detected most microcolonies, microcolony outline recognition is very poor, with the methods primarily recognizing a small region in the center of the cells. Scale bar is 10 µm.

For images acquired by high-throughput microscopy, focal and illumination differences are prevalent within a single experiment. For example, in experiments in which microbes are imaged in a glass-bottom microplate [23], focal-plane variation exists across the plate and even within one well or field. In addition, illumination of the fields in a well’s center is more uniform than in its corners. Illumination unevenness also exists within single imaging fields, both spatially and temporally. It is therefore critical that an effective image-analysis algorithm should be able to: 1) identify objects even under non-optimal focus or brightness, and 2) perform consistent recognition and measurement of the same object across a range of focal planes and illuminations.

To assess the degree to which PIE is robust to this technical noise in colony measurements associated with variability in image illumination and focal plane, we compared the accuracy of microcolony outline recognition (IoU) by PIE and the other cell recognition methods across a range of illumination levels and focal planes of the same imaging position. PIE not only performed consistently in colony identification across both a wide range of focal planes (∼7 µm, equivalent to ∼2 cell diameters) and image brightnesses, but also accurately identified virtually all colonies within the images (**Fig 4**). The other methods displayed lower accuracy in colony recognition than PIE at most focal planes and all exposure times tested.

**Fig 4.**
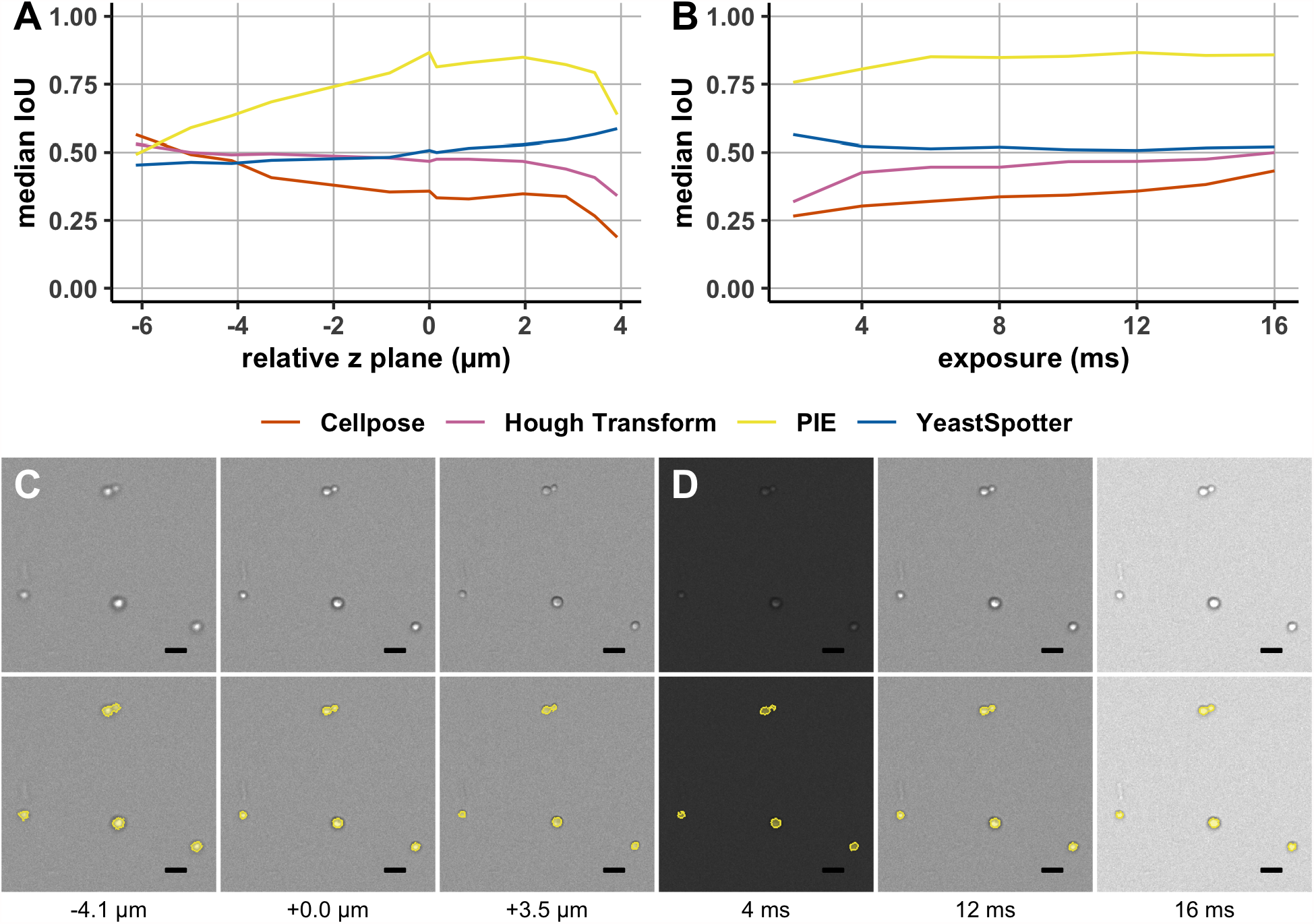
PIE performance across focal plane and illumination conditions. (A-B) Median IoU values across a range of focal planes (*A*) and exposure values (*B)*, with corresponding images of PIE-generated colony outlines for a representative portion of the image tested, (C) and (D) respectively. The top panel of C and D shows the original image, and the bottom panel shows the image with PIE segmentation overlaid. PIE consistently maintains accurate colony outline recognition in highly underexposed or overexposed images. Although the precise outline detected by PIE varies somewhat across focal planes, PIE detects colonies with an IoU of >0.7 across ∼2 cell diameters’ range of focal planes. Note that the middle panel in (C) and (D) (0.0 µm, 12 ms) is identical to the PIE panel in **Fig 3B**, and included only for ease of comparison. Scale bar is 10 µm.

Finally, PIE is significantly faster than alternative methods. We benchmarked average image processing speed on a CPU for PIE against YeastSpotter and CellPose. Average time to analyze a single 5.5-megapixel image on an Intel Platinum 8268 CPU with 8 Gb RAM was 205 seconds using CellPose, 24 seconds using YeastSpotter, and 2 seconds for PIE (13 when PIE was run with cleanup).

We have demonstrated that PIE performs yeast cell recognition more accurately than a ‘naive’ approach with commonly used image analysis software, and both more accurately and significantly faster than two existing deep learning-based approaches trained on high-resolution cell images. PIE-based image recognition is also robust to variation in image brightness and focal plane, making it applicable to real-world high-throughput imaging applications.

### PIE allows population-level measurements of growth properties in diverse microbes

We next assessed the effectiveness of PIE in recognizing growing microcolonies from a wide range of organisms and microscopy platforms. We ran PIE on images of large *S. cerevisiae* microcolonies, as well as either microcolonies or individual cells (for non-colonial microbes) of six additional species: a fungus (*Schizosaccharomyces pombe*), two protists (*Seminavis robusta* and *Trichomonas vaginalis*), a green alga (*Chlorella sorokiniana*) and three bacteria (*Escherichia coli, Pseudomonas aeruginosa*, and *Staphylococcus aureus*) (**Fig 5**). Colony outlines of all seven are effectively recognized, indicating that PIE can be used for automated cell or microcolony recognition in high-throughput experiments across a wide range of microbes.

**Figure 5.**
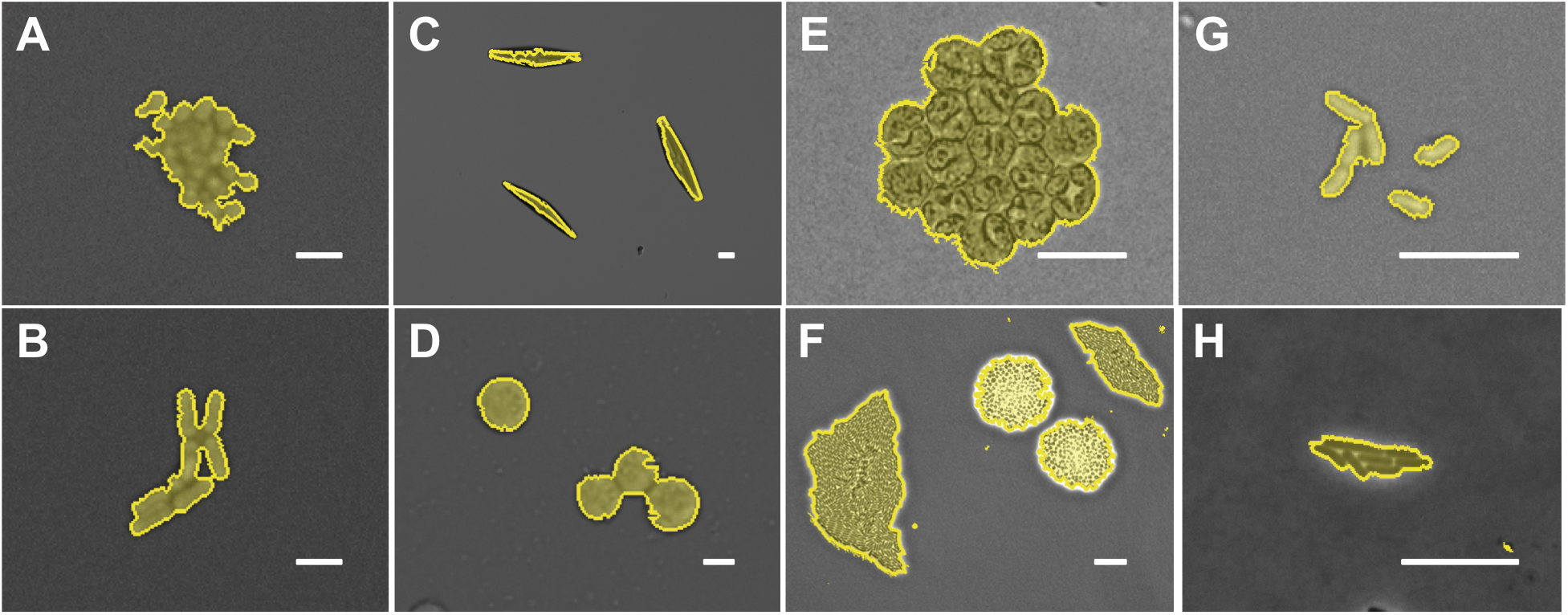
PIE successfully recognizes cell or microcolony outlines across a wide range of microbes. (A-B) Brightfield images of *S. cerevisiae* and *S. pombe* microcolonies, respectively. (C-D) Brightfield images of individuals of *S. robusta and T. vaginalis*, respectively (E) Brightfield image of *Chlorella sorokiniana* microcolony (F) Phase-contrast image of co-cultured microcolonies of *P. aeruginosa* and *S. aureus* microcolonies. (G-H) Brightfield and phase-contrast images, respectively, of *E. coli* cells and microcolony. Scale bar is 10 µm. Images in (C) from Luke Noble and Matthew Rockman; (D) from Sally Warring and Jane Carlton; (E) from Dietrich Kohlheyer [24]; (F) from Dominique Limoli and Kaitlin Yarrington [25] (H) from Jeffrey Carey and Mark Goulian [26], used with authors’ permission.

Robust automated recognition of microcolonies allows large-scale quantification of microbial growth properties by tracking many colonies across a timeseries. We therefore applied PIE to time-lapse microbial imaging to estimate microbial growth rates. PIE calculates the area of each tracked colony and then uses the change in colony area over time to calculate growth rate and pre-growth lag time (**Fig 6A**,**C**,**E**). We imaged tens of thousands of clonal *S. cerevisiae* (**Fig 6B**), *S. pombe* (**Fig 6D**), and *E. coli* (**Fig 6F**) microcolonies founded by single cells and grown in media while adhered to the bottom of a 96-well microscope plate over the course of 10-15 hours, and analyzed the resulting images with PIE. After filtration (see **Software Implementation**), PIE returned growth rates and pre-growth lag values for ∼81,000 *S. cerevisiae* colonies, ∼17,000 *S. pombe* colonies, and ∼46,000 *E. coli* colonies (the difference in sample sizes is primarily the result of differences in cell plating density and the number of imaged fields between the three experiments, and not an organism-specific feature of the assay). In defined SC media, *S. cerevisiae* grows faster than *S. pombe* (median growth rates of 0.40/hr and 0.22/hr, respectively), although both experienced similar lag times (2.1 hrs and 2.2 hrs); the difference in growth rates is likely related to the fact that the SC media used was not optimal for *S. pombe*. As expected, *E. coli* grow faster than either yeast species, with a median growth rate of 0.74/hr and a median lag of 1.0 hrs (of note, we included a PIE filtration requirement for this analysis to remove colonies whose R^2^ for growth rate fit was below 0.9; for *E. coli*, this resulted in the removal of ∼26,000 colonies with growth rates very close to 0).

**Figure 6.**
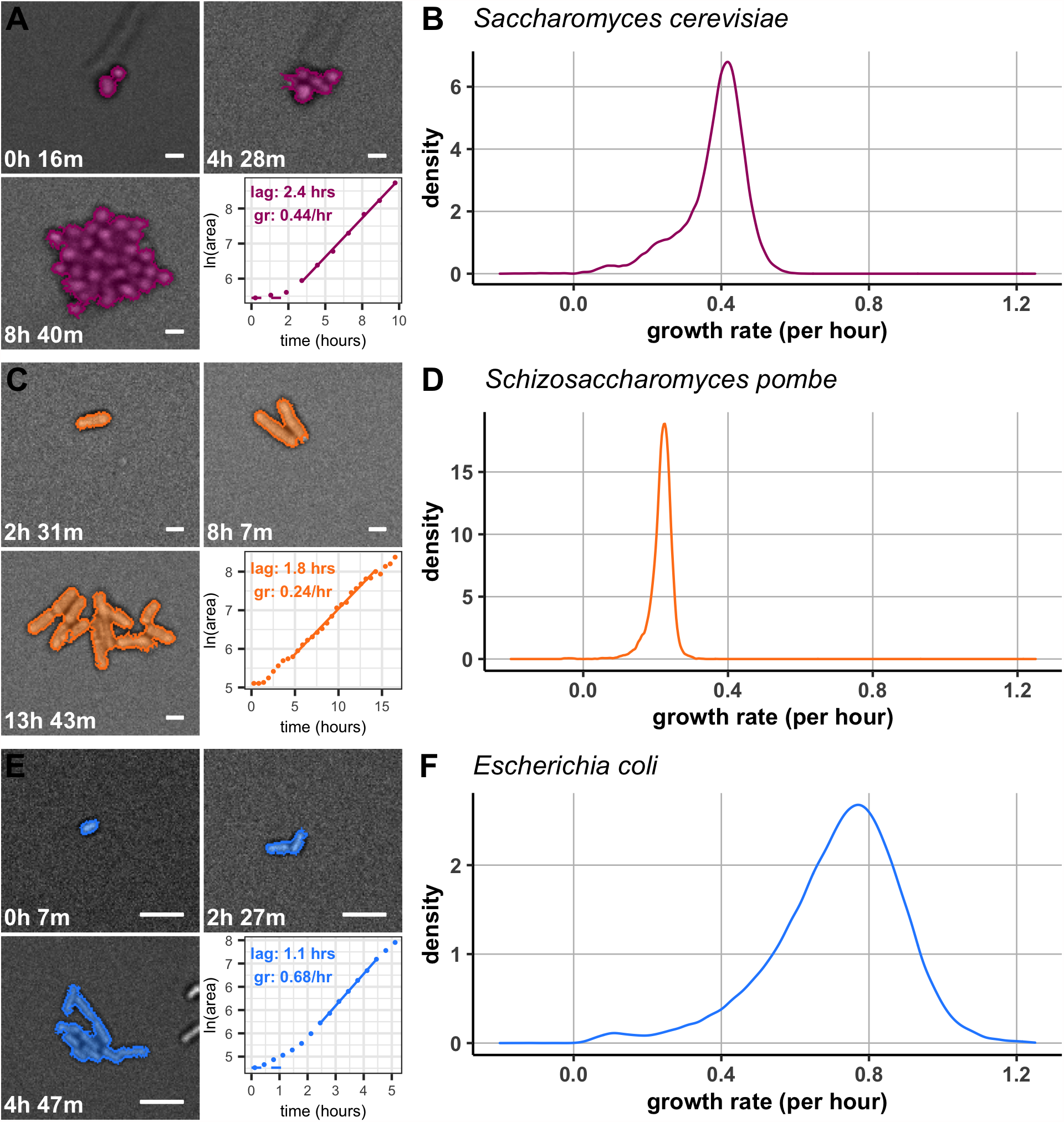
Distributions of individual microcolony growth rates in large populations. Growth of a single representative microcolony from each species, and distribution of growth rates of individual microcolonies of (A, B) *S. cerevisiae*, n = 80,753, (C,D) *S. pombe*, n = 16,625, and (E,F) *E. coli*, n = 46,087. In panels on the left, colony area as determined by PIE is shaded, and the change in *ln(area)* over time used to calculate growth rate and lag is shown in the inset graph, with solid and dotted lines representing best fits for growth rate and lag, respectively. Scale bar is 5 µm.

In addition to generating precise average growth rates, the individual microcolony growth parameters reported by PIE allow exploration of full distributions of growth rates. The three microbe species differ not only in their median growth rates but also in the shapes of their growth-rate distributions. As previously reported [2,7], we found that the distribution of growth rates of *S. cerevisiae* is left skewed (skewness = −1.3), with a small subpopulation of slow-growing cells. We observed a similar skew in the microcolony growth rates of *S. pombe* (skewness = −2.1) and, to a smaller degree, *E. coli* (skewness = −0.7). However, the distributions of microcolony growth rates of *S. cerevisiae* and *E. coli* were much more dispersed than that of *S. pombe* (CV of 0.24, 0.15, and 0.25 for *S. cerevisiae, S. pombe*, and *E. coli*, respectively). These observations reveal the utility of PIE in exploring the non-genetic differences among individual microbes within a population.

### Measurement of individual microcolony-level response to stress using PIE

In addition to tracking individual colonies through time, PIE allows data to be collected in multiple ‘phases’, each with its own imaging regime and analysis parameters; colonies are then tracked across these phases, with growth properties calculated separately for each phase. The effects of perturbations on the growth of individual microcolonies can therefore be tracked, and juxtaposed with each colony’s pre-perturbation properties.

To assess PIE’s utility in tracking multiple colonies before and after an experimental perturbation, we performed an experiment to assess the effect of UV stress on yeast growth. Mutations in the gene encoding histone H2A variant Htz1 have previously been shown to sensitize yeast to UV stress [27]. Furthermore, *htz1-* deletion strains have higher variance of growth rates, and thus a higher proportion of slow-growing cells, than wild type yeast [2]. Slow-growing members of clonal yeast populations have been shown to be resistant to heat shock stress [2,7]. We therefore sought to use high-throughput imaging in combination with PIE to investigate whether (1) there is a relationship between growth rate and resistance to UV stress, and (2) whether the UV stress response is modulated by Htz1.

As discussed in **Software Implementation**, PIE can use fluorescence data to classify colonies co-cultured in a single well. A reference strain can therefore be included in each well of an experiment, increasing statistical power for detecting differences between test strain(s) and the reference. Here, we used a strain expressing a transgene encoding GFP-tagged histone H2B, *Htb2-GFP*, as the wildtype in-well control strain to which *htz1-* colonies were compared. The strains were grown together for ∼3.5 hours while being imaged in brightfield, then a single image in the GFP channel was collected at each position for colony classification (∼40 minutes), and the cells were subjected to a 45-minute UV stress. Following the UV stress, the cells were placed back on the microscope and brightfield imaging continued for ∼9 hours (**Fig 7A**).

**Figure 7.**
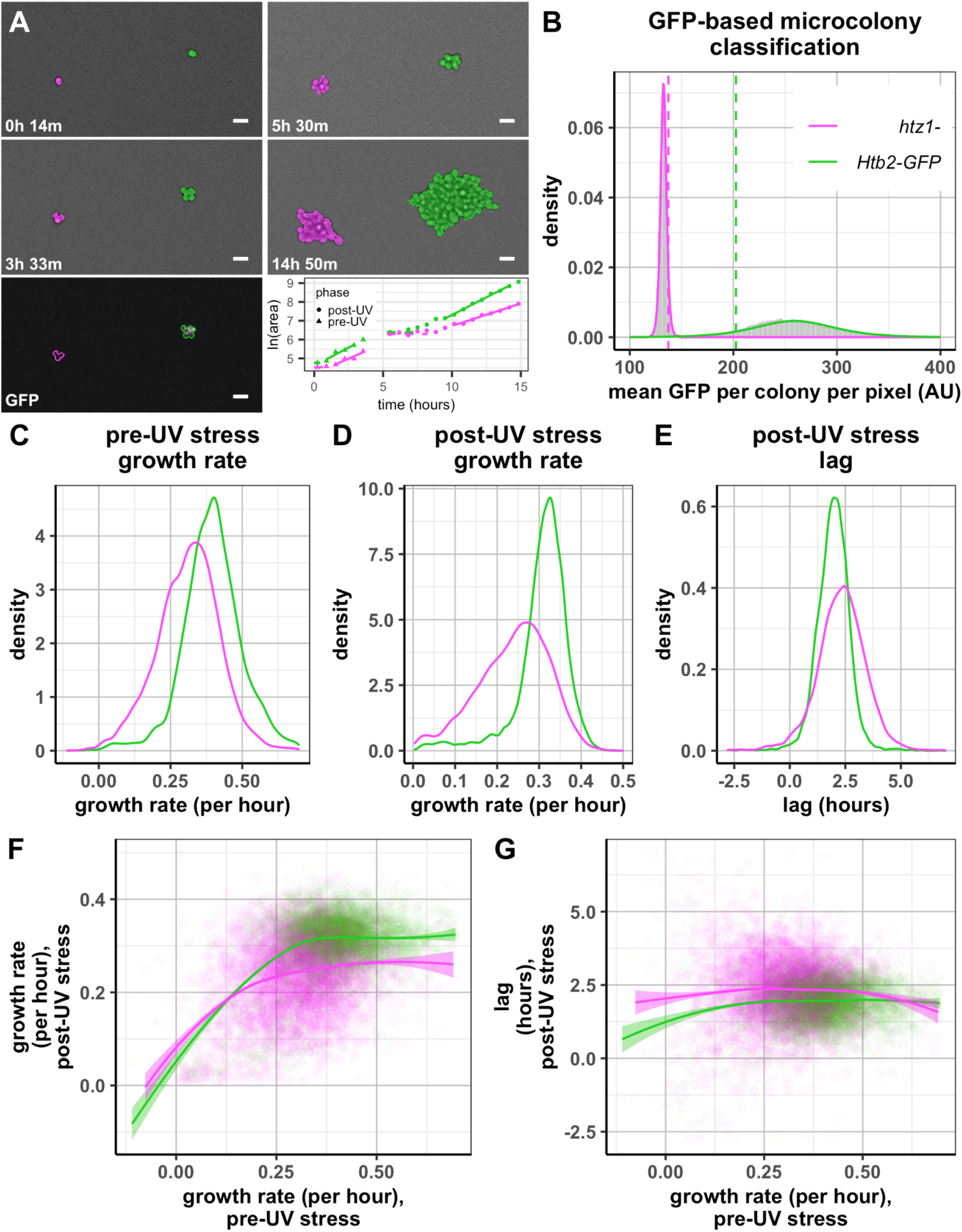
Using multi-phase PIE analysis to study stress response in microcolonies of multiple co-cultured strains. (A) Growth of two colonies over time; left panels are before UV stress, right panels after UV stress. The bottom left panel shows GFP expression (white) overlaid with colony outlines from the previous brightfield image, which are used to calculate per-colony GFP channel pixel intensity. The bottom right panel shows log(colony area) over the two phases of the experiment, with solid and dotted lines representing best fits for growth rate and lag, respectively. The break in datapoints ∼3.5-5 hrs into the experiment was due to collection of GFP fluorescence data and a 45-minute UV treatment; circles denote ln(area) in the pre-stress experiment phase, and triangles in the post-stress experiment phase. Scale bar in images is 10 µm. (B) Histogram (in gray) of mean per-pixel colony GFP fluorescence values, with fitted logistic probability density functions to GFP-negative (*htz1-*, magenta) and GFP-positive (HTB2-GFP, green) colonies, n ∼ 51,000 colonies. Dotted lines represent threshold values used for calling each GFP category; colonies between the thresholds were not assigned to either category. (C-E) Distributions of colony growth rates before and after UV stress, and lag times after UV stress. *htz1*-colonies grow slower than HTB2-GFP colonies both before and after stress, and have a longer post-stress lag period. (F-G) Growth rate and lag after UV stress as a function of each colony’s pre-stress growth rate, with lines denoting the LOESS regression for each strain. n ∼ 12,000 colonies.

To test the ability of PIE to discriminate between the GFP-marked strain and the non-fluorescent *htz1-* strain, we performed our experiment both with wells in which the two strains were co-cultured, and with wells in which only a single one of the strains was present. The results of PIE’s GFP classification can be seen in **Fig 7B** and **Table 1**. To classify colonies, PIE uses data from all wells to identify two thresholds, an upper threshold for the fluorescence-negative colonies, and a lower threshold for the fluorescence-positive colonies (see **Software Implementation**). By using colonies from wells with a single known strain, we can assess the specificity and sensitivity of PIE (**Table 1**). We find that 15%-17% colonies are not classified (i.e. they fall between the thresholds), and only a fraction of a percent of colonies (0.4% for *Htb2-GFP*, 0.1% for *htz1-*) are misclassified into the wrong fluorescence category. The vast majority of colonies (>83%) are classified as their true fluorescence category. It is important to note that, whereas the proportion of misclassified colonies is hardwired in PIE and unlikely to deviate strongly from the values observed here, the proportion of unclassified colonies is highly dependent on the properties of the colony fluorescence distribution, with the lowest percent of unclassified colonies in pairs of strains whose fluorescence levels are maximally separable.

**Table 1.** GFP-based classification of colonies of a known genotype. For each genotype, 15-17% cells were not classified, and < 0.5% of cells were misclassified.

| strain | GFP classification | # colonies | % strain |
| --- | --- | --- | --- |
| *Htb2-GFP* | (+) | 9583 | 84.3% |
|  | None | 1739 | 15.3% |
|  | (-) | 48 | 0.4% |
| *htz1-* | (+) | 12 | 0.1% |
|  | None | 2196 | 16.7% |
|  | (-) | 10951 | 83.2% |

We next examined the pre-and post-UV stress microcolony growth rates of both genotypes, as well as the post-UV stress lag times (**Fig 7C-E)**. As previously reported [2], growth rates were slower and more variable in *htz1*-mutants. UV stress resulted in a protracted lag period, during which colonies did not grow: ∼1.9 hours post-stress lag for *Htb2-GFP* and ∼2.3 hours for *htz1-* colonies. When UV-stressed colonies started growing, growth rates of both genotypes decreased: 22.3% (95% CI: 21.6%-23.0%) in *Htb2-GFP*, and 23.4% (22.5%-24.3%) in *htz1-*. By contrast, in colonies that experienced a mock stress, growth rates increased by a small amount: 3.0% for *Htb2-GFP* (95% CI: 2.3%-3.7%), and 10.8% (10.0%-11.6%) for *htz1-*. The small increase in the mock condition is likely an artifact of the fact that in this experiment, we chose to perform an extended growth phase after UV stress. Because the time window with the maximum growth rate is selected to report the growth rate by our algorithm, the reported growth rate can only increase if additional time points are added. As a result, the absolute growth rate measurement in the longer second stage may appear elevated. This dependence of estimated growth rate on the number of time points is not an issue when relative growth rates of colonies during the same growth phase are compared, but should be considered if absolute growth rates are to be compared across experimental phases. Lag time after mock stress treatment was minimal (0.2 and 0.5 hours for *Htb2-GFP* and *htz1-*, respectively) (**S4 Fig**); the presence of a short lag phase after mock treatment might be the result of a temporary drop in temperature (from 30 °C to room temperature) over the course of the treatment.

In both yeast and bacteria, the growth rate of an individual colony can affect its tolerance of stress [2,7,28]. Because PIE allows colonies to be tracked across multiple phases of a single experiment, we can ask how the growth rates of colonies before stress correlated with the post-stress lag and growth rates of those same colonies (**Fig 7F-G**). Although growth rates of slow-growing cells appear to be positively correlated between the pre- and post-stress phase, growth rates of medium- and fast-growing cells are uncorrelated between phases; these relationships hold for both the *htz1-* and *Htb2-GFP* strains (**Fig 7F**). Post-stress lag also appears uncorrelated with pre-stress growth in medium- and fast-growing cells; most stressed cells that quickly exit lag after stress are found among slow-growing HTB2-GFP colonies (**Fig 7G**). Although these results do not suggest that slow growth confers a strong protective effect against UV stress — as it does against acute heat stress [2,7] — they do suggest that growth heterogeneity contributes to heterogeneity in response to UV stress, particularly in the exit from UV-induced growth arrest.

## Discussion

Here we describe PIE, software that can perform microcolony growth-rate measurements on time-lapse brightfield or phase-contrast images and, if needed, simultaneous measurements of colony fluorescence. PIE combines adaptive cell-center recognition and gradient-based edge detection to recognize outlines of microbial colonies in low-resolution brightfield images and track colonies through time. As we have shown, PIE’s colony recognition is robust to variation in image brightness and focal plane. On low-resolution brightfield images, PIE outperforms existing machine learning-based methods of cell segmentation in both speed and accuracy. The software is highly flexible in terms of experimental setup, allowing multiple phases that might include tracking of growing microcolonies, stress administration and cell classification. PIE is available as a Python package, with full documentation currently at https://pie-image.readthedocs.io/, and we have developed an online application to quickly test analysis of individual images or timeseries, which is currently available at pie.hpc.nyu.edu [29]. Users of the web application may choose to share images they upload with us. While our data suggests that this method generalizes well to a wide range of microbes imaged across diverse conditions, we hope that images shared with us via the web application will allow us to continue to improve PIE.

The ability of PIE to accurately segment cells in low-resolution brightfield and phase-contrast images provides significant advantages over existing approaches to cell recognition in microscopy. The use of brightfield images obviates the requirement for engineered, fluorescent reporter-expressing cells. In addition, no single fluorescence channel is required to be set aside for cell recognition. Furthermore, PIE’s ability to accurately track colonies in low-resolution images vastly increases the potential sample size of any experiment. For example, for the *S. cerevisiae* growth rate data described here, we tracked colonies at 1-hr resolution and measured growth rates for >80,000 colonies in a single experiment. Such sample sizes would not be possible using algorithms that require high resolution (i.e. higher magnification) images.

PIE also does not require any user input into tunable parameters in the colony recognition portion of the algorithm, although advanced users can choose to modify certain default features, e.g. by running the algorithm in ‘cleanup’ mode. In images analyzed for this paper, only three colony recognition parameters were adjusted: whether cleanup was performed, the maximum size of the empty spaces between cells that were considered to belong to the colony, and (for the *P. aeruginosa*/*S. aureus* coculture data) the number of distinct cell body intensities in the image. We include guidelines on all optional image analysis parameters, as well as optional filters used during growth rate calculation, in the PIE documentation. For the latter, the single required filtration parameter is the number of time points in the window used to calculate growth rate. It is designed to allow PIE to select a set of time points for growth rate fitting that includes times when the microcolony is growing, but not points when it is stationary. As a result, if this value is set too high, it will require PIE to include lagging time points in the growth calculation, deflating both the lag and growth rate measurements; if it is set too low, growth rate estimates will be noisy and inflated. In addition, users are encouraged to carefully consider filtration of colonies with a low R^2^ of growth rate fit to area over time; care is required as this filter necessarily also removes colonies with growth rates close to 0.

By providing a simple platform for performing automated simultaneous measurements on microbial growth, PIE opens up opportunities to analyze new and intriguing questions at the single-cell level within large populations. Here, we have highlighted how this software can be used to assess the effect of pre-stress growth parameters on stress resistance. Beyond this example case, PIE reports a wealth of other data, including founder cell size and shape, fluorescence measurements, and colony shape, that can be harnessed to investigate the relationships between gene expression, growth, and stress resistance (e.g. [7]). The variabilities of these traits and their relationships within populations would be especially difficult to measure without an automated image-analytical tool.

We hope that PIE’s ease of use, along with its robustness to experimental variation and the very high sample sizes it affords, encourages others, including those without image-analysis expertise, to use PIE to integrate high-throughput image-based growth assays into their work.

## Materials and Methods

### Software development and availability

PIE was developed in Python 3.7 [30], and is available on GitHub (https://github.com/Siegallab/PIE); documentation for PIE is available at https://pie-image.readthedocs.io/ [29].

Image analysis in PIE is implemented primarily using the OpenCV [31] and Numpy [32] software packages. Curve-fitting steps make use of SciPy [33]. Pandas [34] is used for growth rate filtration and output table compilation steps. Pillow [35] and PlotNine [36] are used for visualization. Click [37] is used for the command-line interface.

An online application for PIE is available at pie.hpc.nyu.edu so that users may quickly test PIE’s analysis of their own images (both individually and in time series) [29]. The online application is implemented using Flask [38].

### Microscopy

To test PIE’s ability to identify and measure colonies at various focal planes and illumination levels, brightfield images of growing cells were acquired using a Nikon Ti Eclipse inverted microscope with Perfect Focus System, which automatically corrects drifts and fluctuations in the Z axis during long-term imaging. Yeast images were acquired with a 10x objective and 1.5x magnifier; bacterial images were acquired with a 40x objective and no additional magnifier. Images were acquired with NIS-Elements software with gain set to 4 and saved as 11-bit .*tif* files. Each well of a 96-well microscope plate (clear bottom MatriPlates, 0.17 mm bottom glass thickness, catalog number MGB096-1-2-LG-L, Brooks Life Science Systems) was filled with 400 μL medium and seeded with cells (as previously described in [2,9,23]).

### Colony outline comparison across cell recognition programs

Two major potential sources of image-recognition error were considered: image brightness and focus. The optimal focal plane (corresponding to slightly defocused cells) and exposure are manually determined to correspond to slightly defocused images where cells have a bright center and a dark outline. Two additional sets of images were taken at the same position as the ‘optimal’ image. One set had the optimal focal plane but varied in brightness (exposure time 2–16 ms), which spans more than the typical image brightness variation across one plate. The other set of images had the optimal brightness (exposure time of 12 ms for this experiment) but varied in focal planes (ranging between −6 µm and +4 µm relative to the optimal focal plane), which spans more than the typical range of focal-plane differences within one experiment. PIE code without cleanup was applied to process these images. The same images were also analyzed using YeastSpotter [14], CellPose [15], and a custom ‘naive’ Hough Transform-based analysis.

For CellPose and YeastSpotter, a median filter was applied to all images before analysis to improve cell recognition by these algorithms, as cell recognition on large raw images was very poor prior to filter application. In CellPose, cytoplasmic recognition using the pre-trained model failed to correctly recognize cells, so nuclear recognition using that model was applied instead and yielded better results.

Hough Transform analysis was performed in Fiji [20–22]. Background subtraction was performed with a rolling radius of 10, followed by Maximum Entropy thresholding, automated edge detection, and Hough Transform with a threshold of 0.65 and minimum and maximum radii of 3 and 7 pixels, respectively.

To calculate F_1_ scores and IoU (Intersection over Union) values (**Eqtn 1** and **Eqtn 2** below), masks created by each method were compared to a hand-traced reference colony mask based on the image with the optimal focal plane and exposure. Before IoU and F_1_ calculation, image registration was performed to align each colony mask to the reference mask; this registration was necessary because the positions of the colonies in the image shifted slightly as the image focal plane was shifted. IoU was calculated based on the number of pixels in each colony and overlap of colonies between the masks. Reference colonies with no matching colony in a given method were assigned an IoU value of 0.

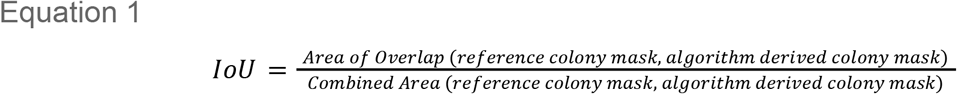

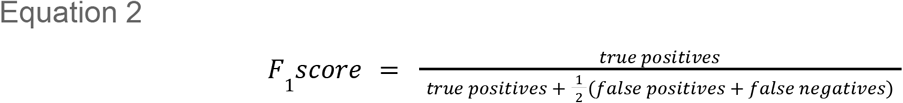

To compare image analysis speed, CellPose, YeastSpotter, and PIE (with and without cleanup) were all run in series on a single Intel Platinum 8268 CPU of a dedicated node on the NYU high-performance cluster, with Turbo mode disabled. For CellPose and PIE, the time to process each individual image was recorded, and values were averaged; for YeastSpotter, images were all processed in a single run, and the process time was divided by the number of images analyzed.

### Strains and Media

For growth rate distribution experiments, *Saccharomyces cerevisiae* of the prototrophic haploid strain FY4 (Winston et al., 1995) were grown to saturation in Synthetic Complete (SC) media, and imaged once per hour in SC media during microscopy. *Schizosaccharomyces pombe* strain 972h-was grown to saturation in Edinburgh Minimal Media (EMM), and imaged once every 30 minutes in SC media during microscopy (because EMM media interferes with ConA immobilization); SC is not optimal for *S. pombe*, and may confer mild stress, but sufficed for the experiments presented here. *Escherichia coli* strain MG1655-F3, a derivative of MG1655 background, containing knockouts of the *fliC* flagellin gene to impair motility and of the *fimA* and *flu* genes to prevent cell aggregation [39] was shared by the Kussell lab. *E. coli* was pre-cultured to saturation in LB media, and imaged every 20 minutes in M9 media during microscopy. For yeast experiments, microscope plates were pre-treated with Concanavalin A (ConA) as previously described [2,23]; *E. coli* were plated on plates in wells treated for 1 hr with 6.6 µL Cell-Tak (Corning, CB402-40) + 53.4 µL 0.1 M sodium bicarbonate, pH 8.0. The demonstration of the successful ‘cleanup’ procedure used haploid strains derived from DBY4974/DBY4975 [40–42]. For the light stress experiment, we used a haploid *htz1-* BY4743 YKO magic marker-derived strain [2,43], and a HTB2-GFP strain from the yeast GFP collection [44].

### Light Stress Experiment

The experiment testing the effects of light stress was performed by growing *HTB2-GFP* and *htz1-* cells to late log phase and plated on ConA-coated plates at 2000 cells/well as described [23]. Wells contained either only one of the two strains or both strains together. Yeast were imaged every 40 mins for 4 hours while growing at 30°C, followed by a single imaging step in the GFP channel to collect fluorescence data. The plate was then immediately placed on a UV transilluminator (Fisherbrand FB-TIV-614A), elevated 3 mm above the transilluminator surface to prevent heating. Half the plate was covered with foil-wrapped cardboard to prevent exposure as a mock stress, and the other half was illuminated with UV on the highest setting for 45 minutes. After the UV treatment, the microplate was placed back on the microscope and the second imaging phase was initiated.

To test the relationship between pre-stress growth rate and post-stress growth rate and lag, we filtered the data, removing the colonies that fell into the upper or lower 0.5% quantile for any growth property (for this analysis, we did not include a filter on minimal R^2^ value of growth rate fit, which results in a very small number of colonies with unrealistically large or small lag/gr values). To model the effects of stress (or mock stress) exposure on each strain, we split data by strain and UV exposure status, and used the *lme4* package in R [45] to fit a linear mixed effect model of growth rate as a function of phase for each subset of the data, accounting for random effects from individual colonies and wells:

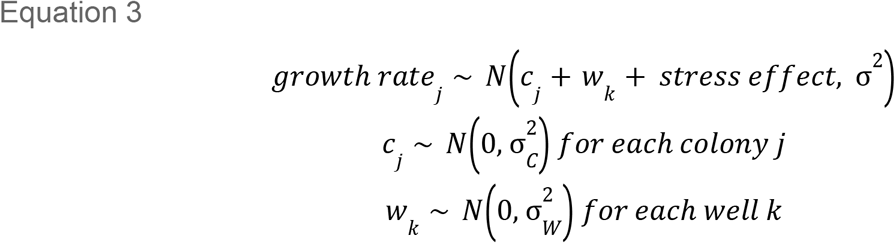

where σ_W_^2^ and σ_C_^2^ are the standard deviation of growth rate across wells and colonies, respectively.

We modeled the post-stress lag in each strain as follows:

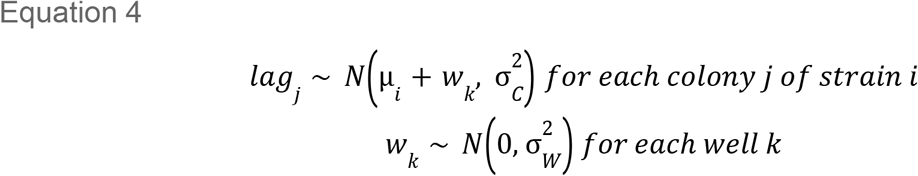

where σ_W_^2^ and σ_C_^2^ are the standard deviation of post-stress lag across wells and colonies, respectively, and µ_i_ is the mean post-stress lag of strain *i*.

## Data availability

Data and code used in this project is available at https://osf.io/6g9n5/?view_only=eb963022aa51416e9dbe0893d04905ef

## Acknowledgments

We thank Luke Noble, Matthew Rockman, Sally Warring, Jane Carlton, Kaitlin Yarrington, Dominique Limoli, Bastian Wollenhaupt, Dietrich Kohlheyer, Jeffrey Carey, and Mark Goulian for sharing microbial images; Edo Kussell and Dan Pollack for sharing the *E. coli* strain used in this study; Fei Li and Josephine Ban for sharing the *S. pombe* strain used in this study; NYU’s Data Science and Software Services and High-Performance Computing, especially Eric Borenstein and Flannon Jackson, for their assistance in setting up the PIE web application and software package; and Shenglong Wang for his extensive help testing and benchmarking PIE throughout the development process.

This work was supported by National Institutes of Health grants R35GM118170 (to MLS) and R01HG008354 (to SFL), the NYU Moore Sloan Data Science Environment (to HM and HJ), and the Google Cloud Research Credits Program. The funders had no role in study design, data collection and analysis, decision to publish, or preparation of the manuscript.

## Supplementary Figures

**S1 Fig.**
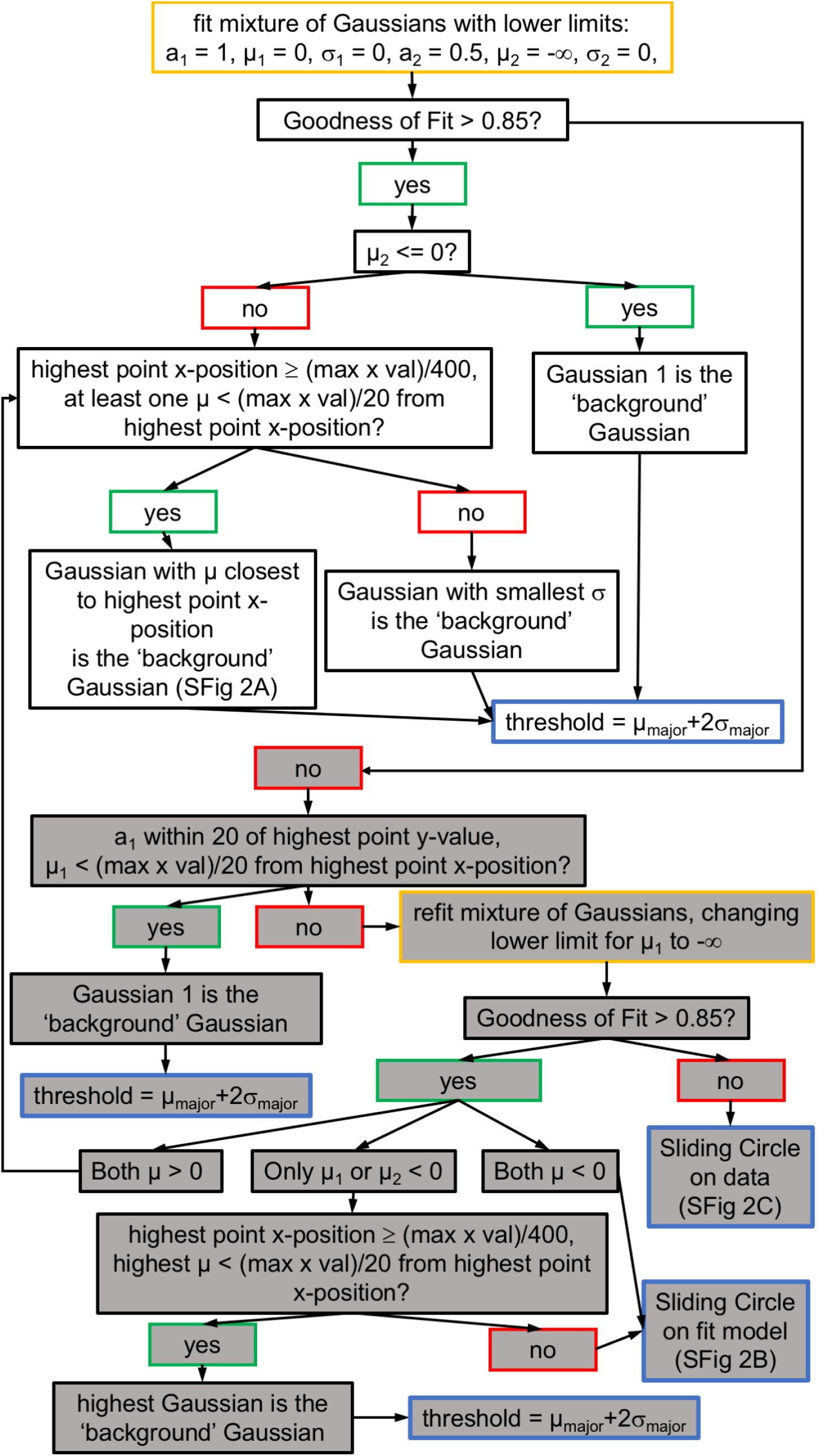
Flowchart of automated threshold identification for cell center recognition. Core threshold-identification algorithm steps are shown with white background; supplemental algorithm steps are shown with gray background. Fit and refit of the data with the two-Gaussian model are denoted with yellow outline. Steps that yield the final threshold are denoted with blue outlines. Example figures for selected steps and all flags for suspect threshold calculation are denoted in parentheses at specific steps. Parameters (with subscript 1 or 2 corresponding to each of the two Gaussians) are: weighting coefficient (amplitude), a; mean, µ; standard deviation, σ. Ideally, both distributions have positive means that are clearly distinguished (**S2A Fig**), but this situation may not always hold. To improve the robustness of automated threshold identification, we developed a supplemental algorithm to which images are passed if they fail to produce an adequate fit to the two-Gaussian mixture model with one mean constrained to be positive (adjusted R^2^ ≤ 0.85). To do this, we considered all possible shapes of the smoothed log-frequency histogram of the top-hat image, and subjected a wide range of problematic images to testing. The supplemental algorithm first tests the source of the poor fit. If the ‘background’ Gaussian distribution is a good fit for the histogram (i.e., the major Gaussian distribution has a similar amplitude and mean to the highest point in the histogram), then the low R^2^ is only caused by a poor fit to pixels corresponding to cell bodies. In this case, the backgroundGaussian distribution’s parameter values are used to determine a threshold, as this Gaussian is a good fit for the background pixels, and this is the only needed part for setting the proper threshold. Otherwise, the two-Gaussian model is refit with no constraints on the means. When this fit is very good (adjusted R^2^ > 0.85), the algorithm attempts to identify which Gaussian corresponds to background pixels by simple comparisons of the two Gaussian distributions’ parameter values. In some cases, neither Gaussian can confidently be called the background-pixel distribution. In those cases, a sliding-circle method described below is applied to the fit model for threshold determination. If the fit to the two-Gaussian model with unconstrained means is inadequate (adjusted R^2^ ≤ 0.85), then the sliding-circle method is applied not to the fit model but to the Savitzky-Golay filter-smoothed data for threshold determination. The sliding-circle method aims to find the nadir of the valley between the two peaks. To do this, it applies a circle mask that glides with its center moving along the smoothed data or fit model. At each position, the ratio of the area within the circle that falls below the curve to that above the curve is calculated. The threshold is determined as the position where this ratio, summed across a given position and two of its neighbours on each side, is maximized. Including neighbour points here reduces the impact of random noise in the pixel-intensity values on threshold determination. Because this sliding-circle method is more time-consuming than the curve-fitting method described above, it is only applied when the two-Gaussian fitting procedures fail. Full details of the fitting and threshold-calculation procedure can be found in the commented code.

**S2 Fig.**
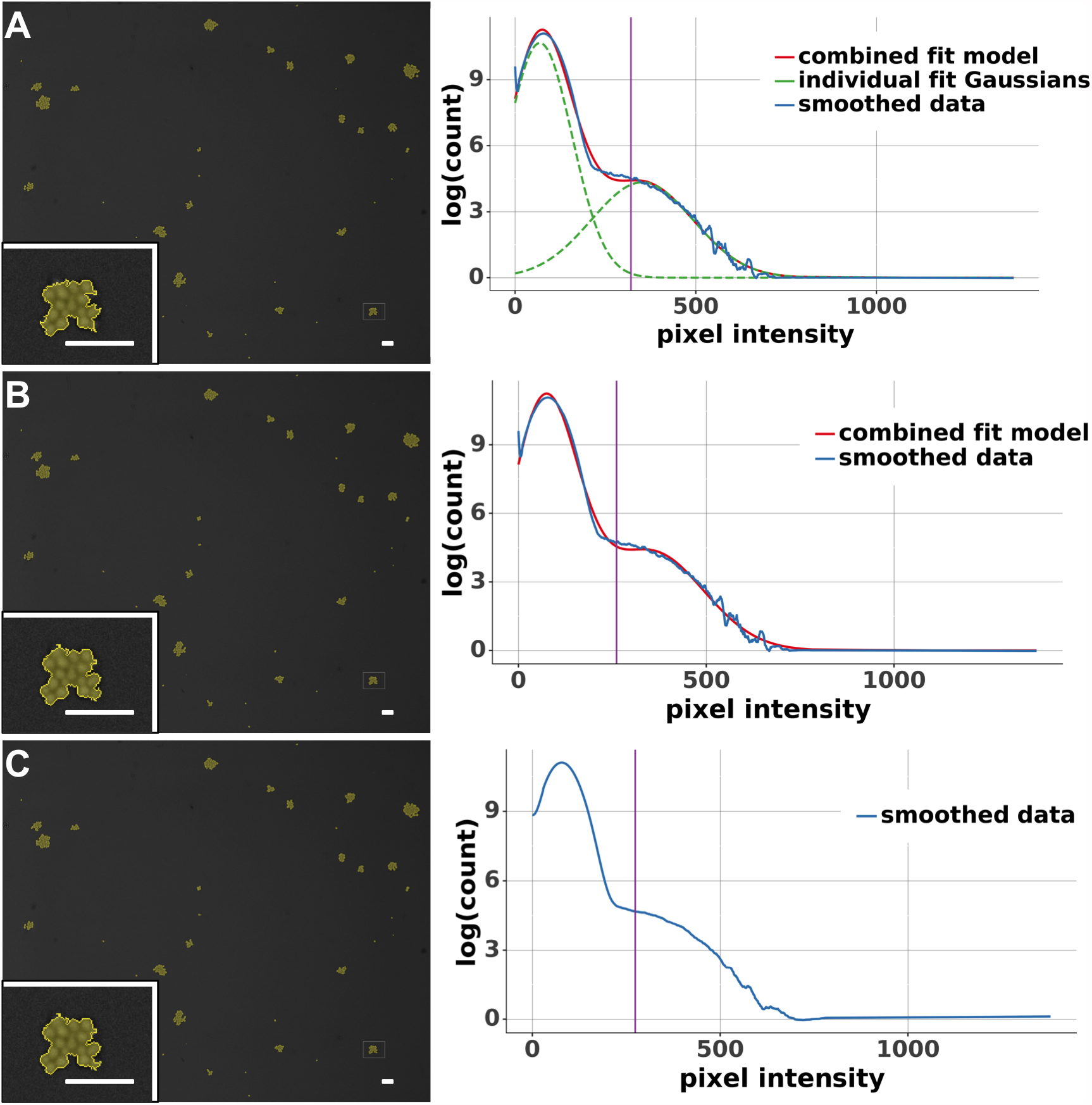
Examples of various approaches to image thresholding. Left panel: Overlays (yellow) of final colony objects recognized by PIE overlaid on a single brightfield image thresholded in different ways. (Inset: Zoomed-in view of the 130×100-pixel region indicated by white rectangle, to show representative colonies and corresponding recognized outlines.) Scale bar = 25 µm. Right panel: Vertical axis is the density of the log count of pixel intensities of the tophat-filtered image. Blue solid curve shows the smoothed log-frequency histogram of the top-hat image; green dashed curves are the two Gaussian fits; red solid curve is the fit combining the two Gaussians; purple vertical line is the calculated threshold for cell centers in the top-hat image. (A) Default two-gaussian thresholding for which both Gaussian fits have positive means and one mean is far away from the highest point of the smoothed data curve; here, the left-most gaussian represents the ‘background’ pixels. (B) Thresholding performed by applying the sliding-circle method to the two-gaussian fit in (A). (C) Thresholding using the sliding-circle method, which is applied to the smoothed log-frequency histogram of the top-hat image directly (note different smoothing window size used here compared to A and B). Note that despite different thresholds identified in A-C, the resulting colony mask is nearly identical, as a result of gradient-based colony edge detection.

**S3 Fig.**
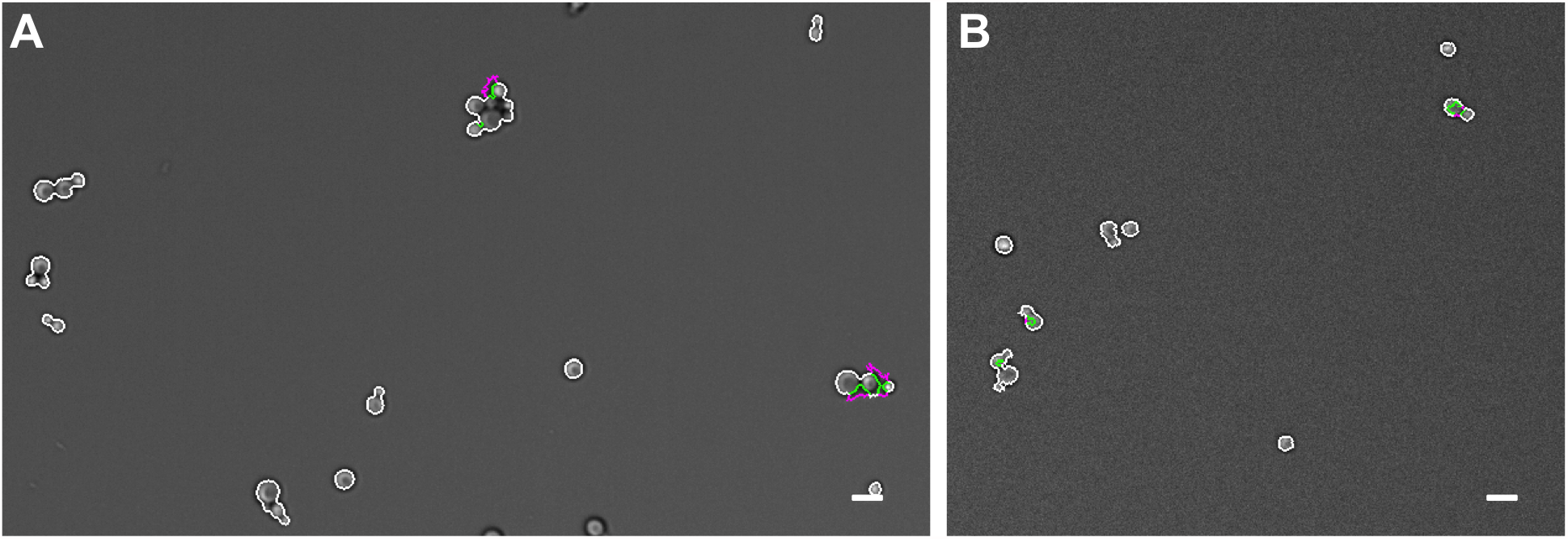
Example images with and without cleanup procedure. (A) An image in which colony outline recognition is improved by the cleanup procedure and (B) an image in which colony outline recognition with the cleanup procedure results in partial loss of colony area from tracking. Magenta, outline without cleanup; green, outline with cleanup; white, overlap. Scale bar = 10 µm.

**S4 Fig.**
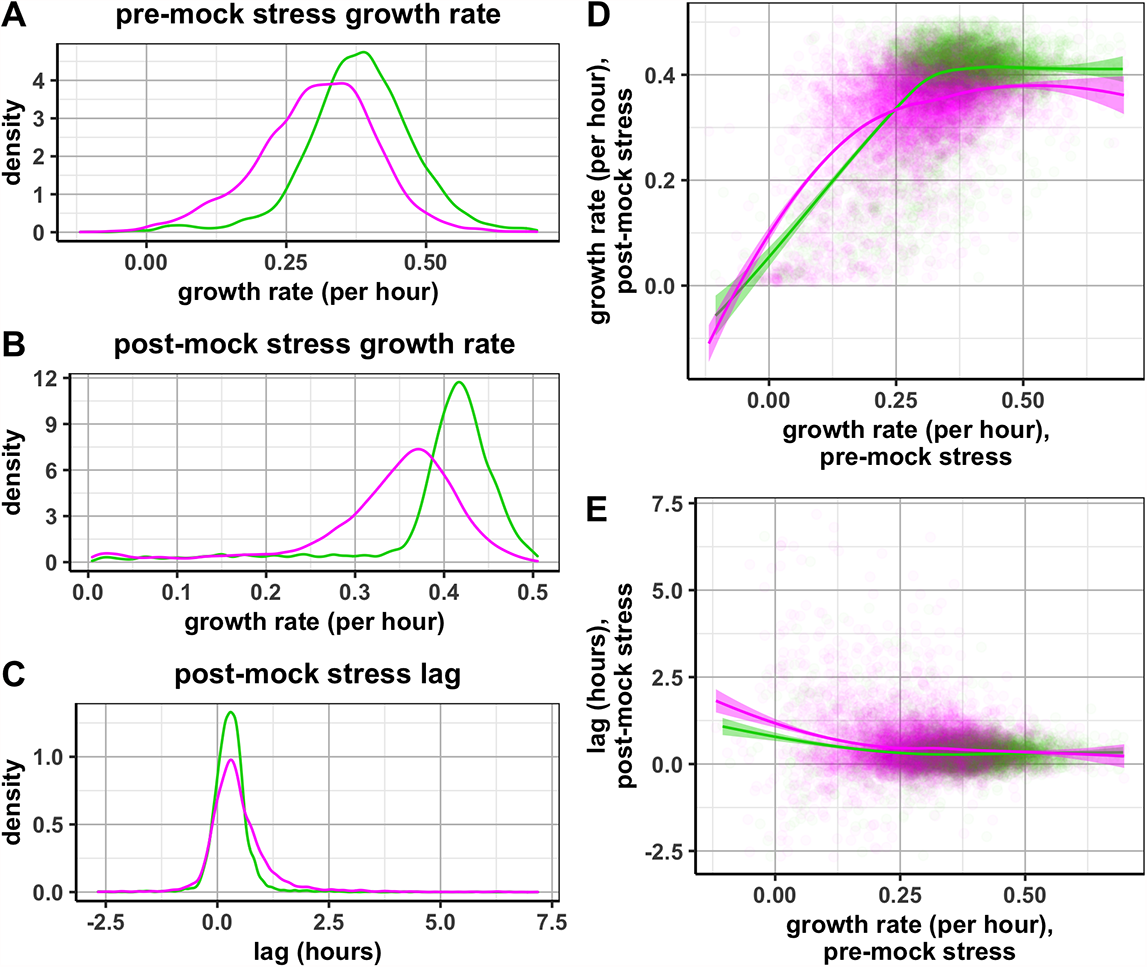
Growth rates and lag times in mock UV-treated yeast. (A-C) Distributions of colony growth rates before and after mock UV stress, and lag times after mock UV stress. *htz1*-colonies grow slower than HTB2-GFP colonies, but the change in mean growth rate after the mock treatment is small, and lag is minimal (D-E) Growth rate and lag after mock UV stress as a function of each colony’s pre-mock stress growth rate, with lines denoting the LOESS regression for each strain. N ∼ 11,000 colonies.

